# Exposure to Traffic-Related Air Pollution is Associated with Greater Bacterial Diversity in the Lower Respiratory Tract of Children

**DOI:** 10.1101/2020.12.09.417501

**Authors:** Christine Niemeier-Walsh, Patrick H Ryan, Jaroslaw Meller, Nicholas J. Ollberding, Atin Adhikari, Tiina Reponen

## Abstract

**Background:** Exposure to particulate matter has been shown to increase the adhesion of bacteria to human airway epithelial cells. However, the impact of traffic-related air pollution (TRAP) on the respiratory microbiome is unknown.

**Methods:** Forty children were recruited through the Cincinnati Childhood Allergy and Air Pollution Study, a longitudinal cohort followed from birth through early adolescence. Saliva and induced sputum were collected at age 14 years. Exposure to TRAP was characterized from birth through the time of sample collection using a previously validated land-use regression model. Sequencing of the bacterial 16S and ITS fungal rRNA genes was performed on sputum and saliva samples. The relative abundance of bacterial taxa and diversity indices were compared in children with exposure to high and low TRAP. We also used multiple linear regression to assess the effect of TRAP exposure, gender, asthma status, and socioeconomic status on the alpha diversity of bacteria in sputum.

**Results:** We observed higher bacterial alpha diversity indices in sputum than in saliva. The diversity indices for bacteria were greater in the high TRAP exposure group than the low exposure group. These differences remained after adjusting for asthma status, gender, and mother’s education. No differences were observed in the fungal microbiome between TRAP exposure groups.

**Conclusion:** Our findings indicate that exposure to TRAP in early childhood and adolescence may be associated with greater bacterial diversity in the lower respiratory tract. Asthma status does not appear to confound the observed differences in diversity. It is still unknown whether the development of asthma changes the lower respiratory tract microbiome or if an altered microbiome mediates a change in disease status. However, these results demonstrate that there may be a TRAP-exposure related change in the lower respiratory microbiota that is independent of asthma status.

## Introduction

For many years it was believed that the lungs were sterile due to the limitation of characterizing bacterial communities through culture-dependent methods [1]. However, with the advancement of molecular-based microbial identification techniques, numerous studies have confirmed that the lungs, do in fact, contain bacterial communities [2-7], and that the microbial community composition may play a role in the exacerbation of chronic lung disease [8].

Exposure to particulate matter (PM) increases the adhesion of bacteria to human airway epithelial cells and PM-stimulated adhesion is mediated by oxidative stress and the receptor for platelet-activating factors [9]. It is also well documented that exposure to traffic-related air pollution (TRAP), a complex mixture of pollutants including fine and ultrafine PM, is associated with incident and exacerbation of existing asthma [10-20]. Although the component of TRAP responsible for this association has not been identified, studies indicate that carbonaceous PM may have a major role in adverse health effects [21-24]. Ultrafine particles dominate particle number concentrations in outdoor urban air, and carbon particles are a major component of these. These ultrafine particles can agglomerate, be retained by the lung tissue upon inhalation, induce pulmonary oxidative stress, and stimulate proinflammatory cytokine release from airway cells [25-28]. There is also evidence that PM promotes airway bacterial infection by weakening the production of an antimicrobial peptide, β-defensin-2 [29]. Several epidemiological studies have documented an association between pneumonia and urban PM [30-32]. Additionally, both the bacterial load and diversity were found to be greater for asthmatic than non-asthmatic adult participants using high-throughput sequencing of sputum samples [33, 34]. Therefore, it is possible that chronic exposure to TRAP may increase the adhesion of microorganisms to the respiratory tract, altering the microbiome of the respiratory tract over time (Fig. 1).

**Fig. 1-.**
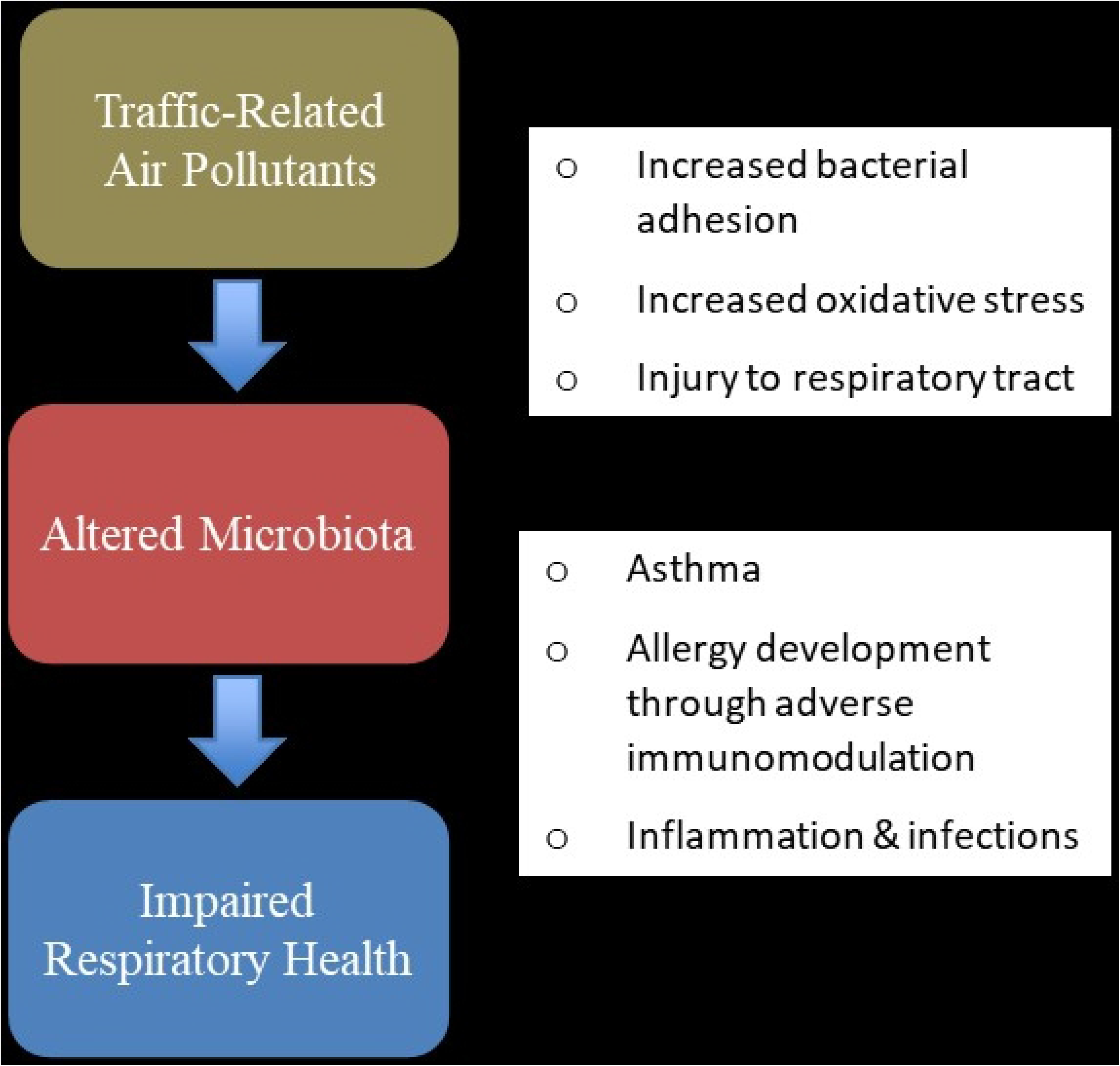
Proposed relationship between traffic-related air pollution (TRAP) exposure and alterations in the lower respiratory tract microbiome leading to impaired respiratory health.

Although recent studies have examined the association of TRAP exposure and the gut microbiome, to our knowledge, there have been no reports on the effects of TRAP on the human respiratory tract microbiome, particularly among children [35-39]. The gut and the lungs are connected via a common mucosal response, as lymphoid cells can travel between mucosal membranes and cause an inflammatory response in multiple areas of the body [40-42]. Therefore, it is plausible that the gut and respiratory system are impacted simultaneously by TRAP exposure. Previously, we studied the association between respiratory end environmental microbiome. We found that the environmental microbiome affected the fungi found in saliva but not bacteria found in saliva or sputum [43].

In this study, we examined the association between childhood exposure to TRAP and the microflora in the lower respiratory tract of children. The participants were recruited from a longitudinal cohort, followed from birth through early adolescence, with a well-characterized TRAP exposure history.

## Methods

### Recruitment

We recruited adolescents enrolled in the Cincinnati Childhood Allergy and Air Pollution Study (CCAAPS) cohort to participate in this study [44]. Briefly, children were enrolled in CCAAPS prior to age one and longitudinally evaluated for allergy and asthma development at clinic visits at ages 1, 2, 3, 4, 7, and 12. Clinic visits included skin prick tests to 15 aeroallergens, spirometry, and questionnaires to assess address history for home, daycare and school, tobacco smoke exposure, pet ownership, home characteristics, and respiratory health [45-49]. At age 12, asthma status was determined by parent report of a physician diagnosis of asthma. Exclusion criteria for this study included having exposure to environmental tobacco smoke at either age 12 CCAAPS clinic visit or at the time of sputum sampling (~ 14 y), or having an upper or lower respiratory infection within 4-weeks prior to sputum sampling, following the protocol by Tunney et al. [50]. We also collected sputum and saliva samples from four pilot participants prior to recruiting the forty full study participants to confirm that we would be able to obtain a sufficient amount of sputum and bacterial DNA from the healthy adolescent population. The study protocol was approved by the University of Cincinnati Institutional Review Board and informed parental consent and participant assent were obtained prior to study participation.

### TRAP exposure estimation

A land-use regression (LUR) model to estimate exposure to a surrogate of TRAP, elemental carbon attributable to traffic (ECAT), has been developed, refined, and validated as part of the CCAAPS [51, 52]. Briefly, an ambient air sampling network, consisting of 27 sites, was operated from 2001-2006 [51], and samples were analyzed for PM2.5 mass concentrations, 39 elements, and elemental and organic carbon [53]. A multivariate receptor model, UNMIX, was used to determine significant sources contributing to PM2.5, including traffic. The contribution to PM2.5 from diesel traffic was assessed using elemental source profiles identified from measurements conducted at cluster sources of trucks and buses [53]. A marker of traffic-related particles specifically related to diesel combustion, the ECAT, was derived for each sampling site and served as the dependent variable for the LUR model. Significant predictor variables in the LUR model included elevation and nearby truck traffic and bus routes [51, 53, 54]. A time-weighted average (TWA) daily exposure of ECAT was calculated for each age of children taking into account the locations where they spent more than eight hours per week, up until age 12, where only the home address was used. For this study, we categorized participants as exposed to high or low levels of TRAP if their average exposure from birth through age 12 was above or below the median exposure of all participants who completed the age 12 study visit.

### Sample collection and preparation

Saliva samples and induced sputum samples were collected from participants with the assistance of the Schubert Research Clinic at Cincinnati Children’s Hospital Medical Center. Immediately prior to sputum induction, participants rinsed their mouths with nuclease-free water, and then 2 mL of saliva was collected using the Norgen Saliva DNA Collection Kit (Norgen BioTek Corp., Thorold, ON, Canada) according to the manufacturer’s instructions. Saliva was collected first so that we could compare the bacterial communities in saliva to those in sputum to ensure that the bacterial communities were distinct and that the sputum samples were not entirely contaminated by the oral microbiome. Next, participants underwent a spirometry assessment and received a dose of albuterol. To induce sputum production, participants breathed in a nebulized hypertonic saline solution for five minutes, then coughed the sputum into a Norgen Sputum DNA Collection Kit (Norgen BioTek Corp., Thorold, ON, Canada). This cycle was repeated up to five times, or until 2 mL of sputum had been collected. After adding the Norgen Collection Kit preservative, which preserves the samples for up to five years at room temperature, sputum and saliva samples were stored at room temperature until DNA extraction.

### DNA isolation

DNA from saliva was extracted using the Norgen Saliva DNA Isolation Kit (Norgen BioTek Corp., Thorold, ON, Canada). DNA from sputum was extracted using the Norgen Sputum DNA Isolation Kit (Norgen BioTek Corp., Thorold, ON, Canada). Sputum was first liquefied using a 100 μg/mL solution of dithiothreitol and incubated at 37°C for 60 minutes. The manufacturer’s protocol was used for DNA isolation of sputum and saliva samples, with one modification: after adding the proteinkinase K and lysozyme, the sample was incubated in an ultrasonic water bath at 65 °C for 30 minutes. This modification was made based on recommendations by Luhung et al. [55] for protocol improvements for DNA isolation of biological aerosol samples with low concentrations of DNA, as we expected low DNA yield from these samples based on our pilot samples.

### Real-Time Quantitative-PCR (qPCR)

To measure the total bacterial DNA present in the samples, the universal primers, UniBacteria_F and UniBacteria_R, and probe, UniBacteria_P1, for the amplification of the 16s bacterial rDNA were used as described by Nadkarni et al. [56]. Extracted DNA from a solution of *Bacillus atrophaeus* with a known concentration of cells was used as the standard. For fungi, the universal fungal primers 5.8Fl and 5.8Rl, and probe, 5.8Pl, for the amplification of the internal transcribed spacer (ITS) regions of fungal rDNA were used [57]. Extracted DNA from a solution of *Aspergillus versicolor* with a known concentration of spores was used as the standard. A set of PCR reaction mixtures were spiked with a known concentration of DNA to test for inhibition. A serial dilution was also included in the well plate as an internal standard to check for pipetting errors. Amplification was performed using the TaqMan system on Applied Biosystems StepOnePlus Real-Time PCR System. All qPCR reactions were replicated three times per sample, and the reported value is the mean of the triplicates. Reagent blanks were also included.

### Metagenomics sequencing

For bacterial sequencing, we chose to amplify the V4 region of 16s bacterial rDNA (primer set 515F-806R) with v2 chemistry because these conditions have documented lower error rates [58, 59]. PCR was carried out by adding 3.5 μL of each forward and reverse primer and 4 μL of Master Mix, containing 0.3 μL 10 mM dNTP, 1.5 μL buffer + MgCl_2_, 0.1 μL FastStart Taq DNA Polymerase, and 2.1 μL nuclease-free water, to 4 μL of DNA extract of each sample. The following thermocycling conditions were used for amplification: 94°C for 60 seconds, 30 cycles of 94°C for 30 seconds, 50°C for 45 seconds, and 72°C for 120 seconds, then 72°C for 300 seconds, and lastly, a 10°C hold. Paired-end sequencing (250 x 2) was performed on the Illumina MiSeq (Illumina Inc., San Diego, CA). PCR amplification and sequencing were performed by the Cincinnati Children’s Hospital and Medical Center DNA Core.

For fungal sequencing, we used the ITS1F-ITS2aR primers for the amplification of the ITS1 region. The ITS fungal rDNA region was selected over the 18s rDNA region because it has been found to be more precise in fungal community analysis [60]. The ITS1 region was selected over the ITS2 region because the pair of primers for the amplification of ITS1 have been found to be more selective, producing fewer non-fungal sequences, in addition to producing a higher number of sequences, than the primer pair for the amplification of the ITS2 region [61]. Fungal PCR and sequencing was performed by RTL Genomics, a division of Research and Testing Laboratories.

Primer sequences were removed using cutadapt v1.16 [62]. The open-source R package, DADA2 v1.8, was used to process reads and for error correction [63]. The default parameters were used for quality filtering, error modeling, dereplication, denoising, and merging of paired-end reads. For bacteria, forward reads were truncated at 210 and reverse reads were truncated at 160 nucleotides. For fungi, reads with less than 50 base pairs were removed to get rid of suspicious low-length reads. Reads with a quality score less than or equal to two, with a maximum expected error rate for the forward or reverse read greater than two, or with a forward or reverse read that contained an ambiguous base were removed. After error correction, the forward and reverse reads were merged to form an amplicon sequence variant (ASV) table. The DADA2 function removeBimeraDenovo was used for chimera removal. Silva version 132 was used as the reference database for bacterial taxonomic classification [64]. For fungi, the UNITE ITS database was used as the reference database [65]. Sequences were aligned using the AlignSeqs function in the DECIPHER package v2.12.0 [66]. For bacteria, a de novo phylogenetic tree was generated using the phanghorn package v2.5.5 [67]. For fungi, a de novo tree was created using agglomerative clustering of the sequence distance matrix using USEARCH [68]. Phyloseq v1.28 was used to integrate the sample metadata, ASV table, phylogenetic tree, and taxonomic assignments for statistical analyses [69]. This method was selected over the construction of operational taxonomic units (OTUs) as it has been argued that using ASVs as the unit for marker-gene analysis improves reusability, reproducibility, and comprehensiveness of data [70].

### Data analysis

We accounted for differences in sequencing depth by multiplying the relative abundance values by the qPCR values for each sample to calculate absolute abundance [71, 72]. Two sputum samples were removed from the bacterial dataset due to a low number of reads (<5000 total reads). Four other sputum samples and four saliva samples were not included in the bacterial dataset because the rDNA did not amplify either during qPCR or during PCR amplification prior to 16s sequencing. Four saliva samples were not included in the bacterial dataset because the rDNA did not amplify during qPCR or during PCR amplification prior to 16s sequencing. For fungi, saliva samples were not included if they did not amplify or had <5000 reads after sequencing. Due to low fungal abundance in sputum, sputum samples were included if they amplified and had >400 reads. These samples were treated as pilot samples. The number of observed ASVs and Shannon alpha diversity were calculated using phyloseq::estimate_richness and Faith’s phylogenetic diversity was calculated using the function pd in the picante R package (v1.8.0) [73].

Alpha diversity indices between sample types, between exposure groups, between genders, and between asthma status groups were first univariately compared using the Wilcoxon rank sum test. We used multiple linear regression to model the effect of TRAP, adjusted for gender, asthma status, and mother’s education as a measure of socioeconomic status, on the alpha diversity in sputum. TRAP exposure was modeled both as a categorical (high/low) and continuous (ECAT) variable (high/low) in separate models. The overall bacterial abundance from qPCR was compared between TRAP exposure groups, asthma status groups, and genders using the Wilcoxon rank sum test.

The Adonis function in the vegan package (2.5-6) was used to implement a permutational multivariate analysis of variance to test for differences in beta diversity between sample types. A dispersion test for the homogeneity of variance across sample type was also conducted using the vegan::betadisper and vegan::permutest functions, as differences in variance may confound the Adonis test. Negative binomial regression as implemented by DESeq2 (version 1.24.0) was used to estimate the fold-change of taxa in sputum according to asthma status, gender, and TRAP exposure groups [74]. Taxa that were not observed in at least 20% of samples were excluded from the DESeq2 analyses. The Human Microbiome Project (HMP) R package (v.2.0.1) function xdc.sevsample was used to assess the distribution of major phyla between sample type, and sputum between asthma status, gender, and TRAP exposure groups [75].

## Results

### Characteristics of study participants

#### Bacteria

Sputum samples from 34 of the participants were included in the bacterial analyses. There were 17 participants in each TRAP exposure group for bacteria. The median ECAT in the high exposure group was 0.46 μg/m^3^, and the median ECAT in the low exposure group was 0.29 μg/m^3^. The cut-off between the low and high exposure groups was ECAT = 0.33 μg/m^3^. In the high exposure group, 42% of the participants were female, 35% were asthmatic, and 82% of the mothers had education beyond high school. In the low exposure group, 53% were female, 18% were asthmatic, and 100% of the mothers had education beyond high school.

Saliva samples from 36 participants were included in the bacterial analyses. For saliva samples, there were 16 participants in the low exposure group and 20 participants in the high exposure group. Of the saliva samples included in the high exposure group, 50% of participants were female, 35% were asthmatic, and 85% of the mothers had education beyond high school. Of the saliva samples included in the low exposure group, 50% participants were female, 13% were asthmatic, and 100% of the mothers had education beyond high school.

#### Fungi

Sputum samples from 10 of the participants were included in the fungal analyses. For sputum, 5 participants were in each TRAP exposure group. Of the 10 samples, 60% of participants were female, 50% were asthmatic, and 90% had mothers with education beyond high school. Saliva samples from 8 of the participants were included in the fungal results. For saliva, 4 participants were in each TRAP exposure group. Of the 8 saliva samples, 50% of the participants were female, 37.5% were asthmatic, and 100% had mothers with education beyond high school.

As so few samples returned amplified fungal DNA sequence reads, the fungal results discussed are considered pilot samples.

### Distinction between sputum and saliva

Sputum had a greater median bacterial alpha diversity than saliva for all three diversity measures (Fig. 2). The median Shannon diversity index was 3.7 in sputum and 3.4 in saliva (p<0.001). The number of observed ASVs was 168 in sputum and 157 in saliva (p=0.019). The median phylogenetic diversity index was 8.3 in sputum and 7.7 in saliva (p=0.6).

**Fig. 2-.**
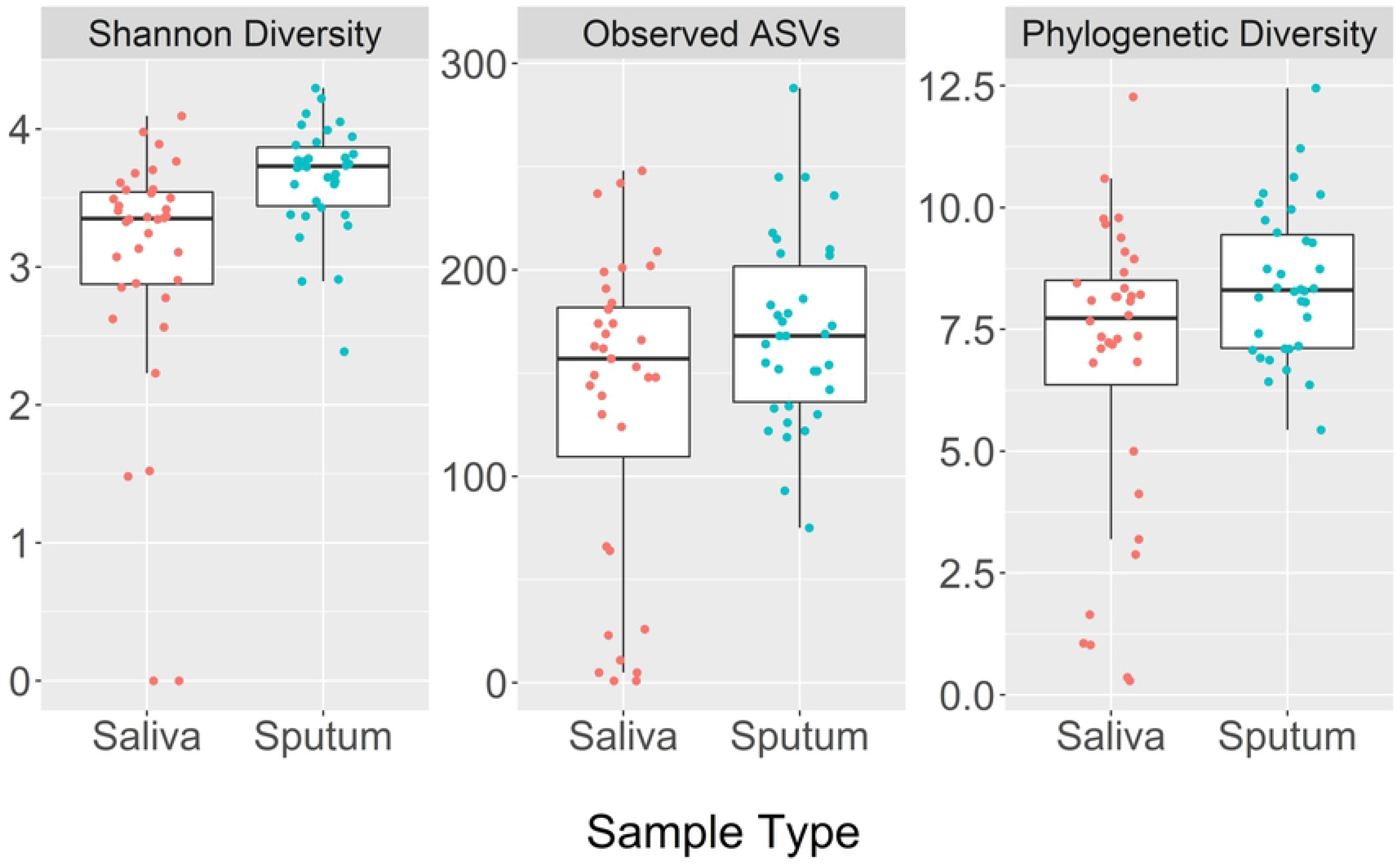
Box plots comparing bacterial alpha diversity indices between sputum and saliva, including Shannon diversity, number of observed amplicon sequence variants (ASVs), and Faith’s phylogenetic diversity.

The most abundant bacterial phyla in both sputum and saliva samples were *Actinobacteria, Firmicutes, Bacteroidetes*, and *Proteobacteria*. All other less abundant phyla were combined into one “other” category for testing the distribution of the phyla between sputum and saliva. The results indicated that the distribution of phyla differed in sputum and saliva (p≤0.001). *Actinobacteria* and *Firmicutes* were more abundant in sputum and *Proteobacteria* and *Bacteroidetes* were more abundant in saliva (S1 Fig).

We used the Bray-Curtis dissimilarity metric to compare the microbial community composition between the two sample types for bacteria (Fig. 3). The Adonis test indicated that 6% of the variance in the distance matrix between sputum and saliva could be attributed to the sample type (p=0.001). The homogeneity of dispersion test failed to reject the null hypothesis that the variances of the sputum and saliva samples were similar (p=0.31) indicating no confounding effect for a difference in variance; however, several outlying saliva samples were observed. We also examined the beta diversity between sputum and saliva using the unweighted UniFrac and Jaccard methods to assess the robustness of the findings to the chosen distance/dissimilarity measure and approach (S2 Fig).

**Fig. 3-.**
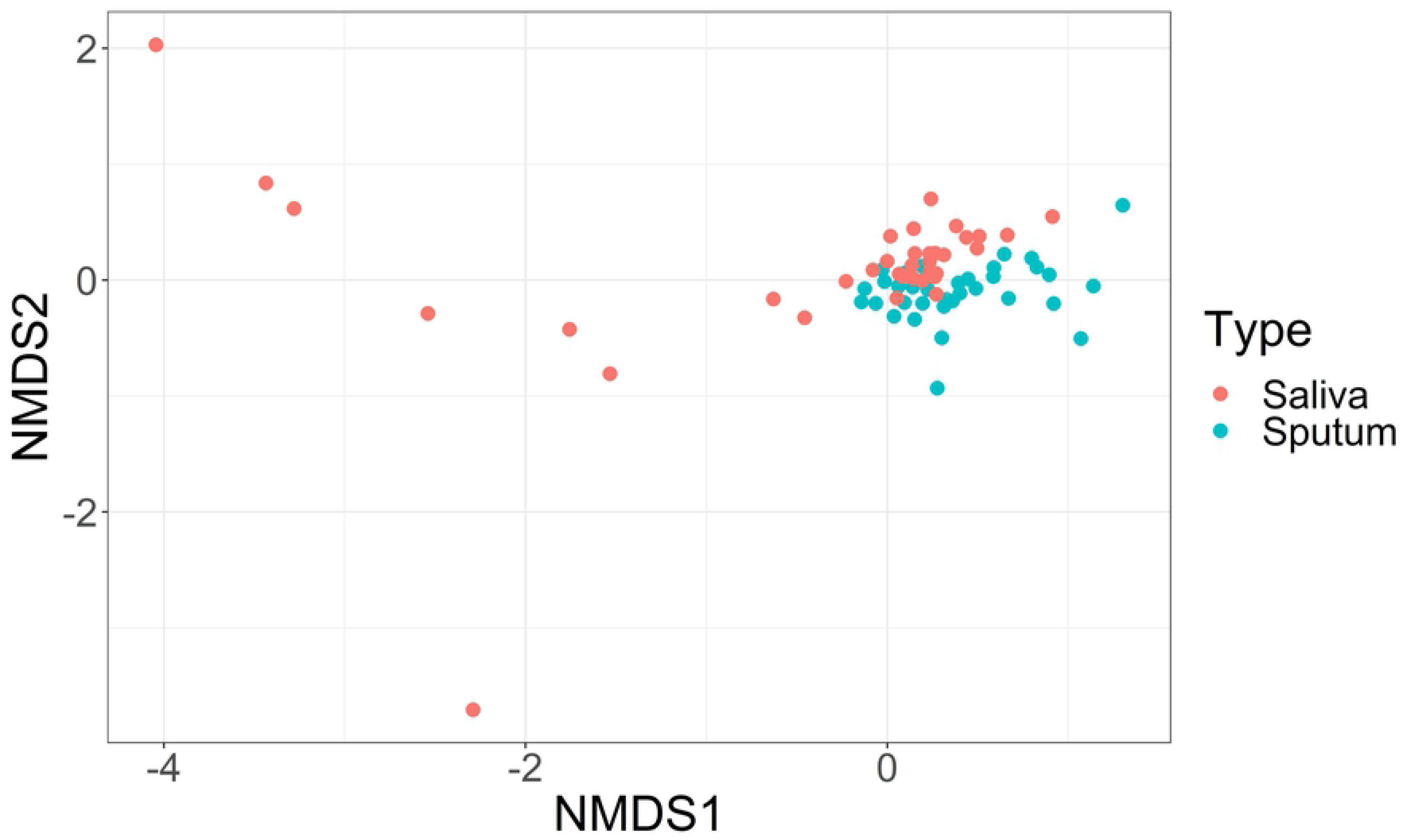
Non-metric multidimensional scaling (NMDS) of Bray-Curtis distances of bacteria in saliva and sputum samples.

### Bacterial Microbiota Differences in TRAP Exposure Groups

#### Alpha Diversity

There was greater diversity in the sputum of the high TRAP exposure group when compared to the low exposure group (Fig. 4). Univariate analysis showed that the number of observed ASVs, Shannon diversity, and Faith’s phylogenetic diversity were all greater in the high TRAP exposure group for bacteria (Table 1). For phylogenetic diversity, there was also a statistically significant difference between genders (Table 1, S3 Fig). There was noticeable within-group variability, suggesting other unidentified factors may be impacting alpha-diversity estimates. There were no statistically significant differences between asthma status groups (Table 1, S3 Fig). In contrast to sputum, there were no observed differences in bacterial alpha diversity in saliva between TRAP exposure group, gender, nor asthma status (S4 Fig).

**Fig. 4-.**
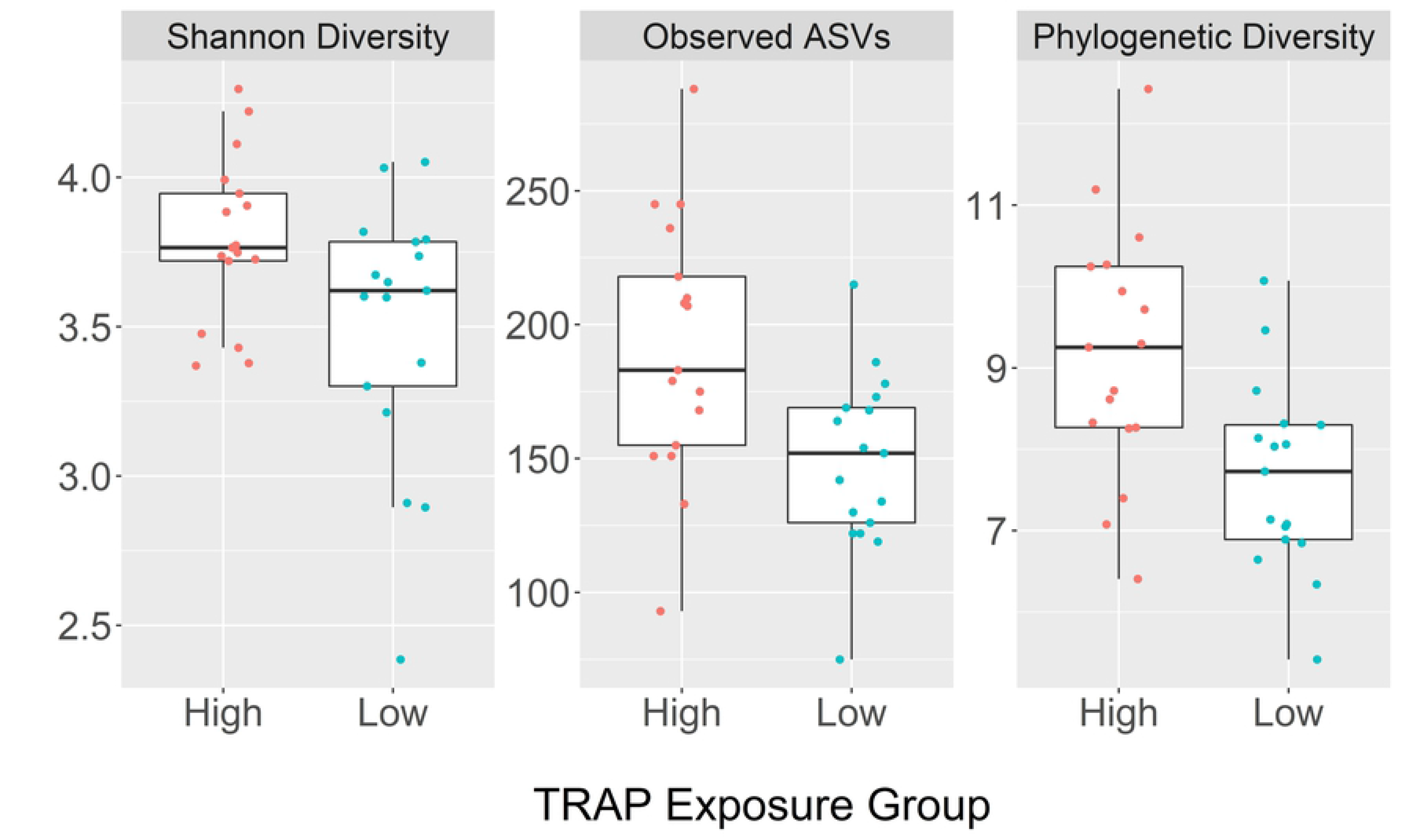
Box plots comparing bacterial diversity indices in sputum between high and low traffic-related air pollution (TRAP) exposure groups; Shannon diversity, number of observed amplicon sequence variants (ASVs), and Faith’s phylogenetic diversity.

**Table 1.** Mean (95% confidence interval) and p-value for each bacterial alpha diversity measure in the sputum by TRAP exposure, asthma status, and gender.

|  | TRAP Exposure |  |  | Asthma Status |  |  | Gender |  |  |
| --- | --- | --- | --- | --- | --- | --- | --- | --- | --- |
|  | High<br>(n=17) | Low<br>(n=17) | p-value | Asthmatic<br>(n=9) | Non-Asthmatic<br>(n=25) | p-value | Female<br>(n=16) | Male<br>(n=18) | p-value |
| **Number of Observed ASVs** | 191 (168-214) | 149 (133-164) | p=0.008** | 179 (154-204) | 166 (147-185) | p=0.38 | 184 (158-210) | 157 (141-173) | p=0.06 |
| **Shannon Diversity** | 3.8 (3.7-3.9) | 3.5 (3.3-3.7) | p=0.05* | 3.7 (3.6-3.8) | 3.6 (3.4-3.7) | p=0.51 | 3.7 (3.5-3.9) | 3.6 (3.5-3.7) | p=0.16 |
| **Phylogenetic Diversity** | 9.2 (8.5-9.9) | 7.7 (7.1-8.3) | p=0.002** | 8.6 (7.6-9.6) | 8.4 (7.8-9.0) | p=0.51 | 9.0 (8.2-9.8) | 7.9 (7.3-8.5) | p=0.04* |

Wilcoxon rank sum test; Significance levels: \*= $p \leq 0.05$ , \*\*= $p \leq 0.01$

Using multiple linear regression, TRAP, both as a categorical (high/low) and continuous (ECAT) variable, was positively associated with an increased number of observed ASVs and phylogenetic diversity after adjusting for asthma status, gender and mother’s education (Table 2). TRAP as a categorical variable was positively associated also with Shannon diversity. Female gender was positively associated with the number of observed ASVs and phylogenetic diversity, but not with Shannon diversity. Neither asthma status nor mother’s education were statistically significant predictors of alpha diversity indices.

**Table 2.** Linear regression model results examining the effect of traffic pollution, asthma status, gender, and socioeconomic status (mother’s education) on the bacterial alpha diversity indices in sputum.

|  |  | Number of Observed ASVs |  | Shannon Diversity |  | Phylogenetic Diversity |  |
| --- | --- | --- | --- | --- | --- | --- | --- |
| | | $\beta$ | p-value | $\beta$ | p-value | $\beta$ | p-value |
| Traffic-related air pollution exposure as a categorical variable (High/Low) | **Intercept** | 214 | **p<0.001\*\*\*** | 3.94 | **p<0.001\*\*\*** | 10.1 | **p<0.001\*\*\*** |
|  | **TRAP Exposure Group (high vs. low)** | 47.6 | **p=0.003\*\*** | 0.32 | **p=0.03\*** | 1.78 | **p<0.001\*\*\*** |
|  | **Asthma Status (yes vs. no vs.)** | 1.15 | p=0.95 | 0.09 | p=0.62 | -0.07 | p=0.91 |
|  | **Gender (female vs. male)** | 34.9 | **p=0.02\*** | 0.12 | p=0.37 | 1.28 | **p=0.009\*\*** |
|  | **Mother's Education (Beyond HS vs. not beyond HS)** | 1.68 | p=0.96 | 0.06 | p=0.84 | 0.53 | p=0.63 |
|  |  | **R<sup>2</sup>=0.36** |  | **R<sup>2</sup>=0.18** |  | **R<sup>2</sup>=0.41** |  |
| Traffic-related air pollution exposure as a continuous variable (ECAT) | **Intercept** | 113 | **p=0.002\*\*** | 3.40 | **p<0.001\*\*\*** | 6.05 | **p<0.001\*\*\*** |
|  | **ECAT** | 202 | **p=0.01\*\*** | 1.00 | p=0.19 | 8.32 | **p=0.002\*\*** |
|  | **Asthma Status (yes vs. no)** | -2.72 | p=0.89 | 0.07 | p=0.70 | -0.24 | p=0.70 |
|  | **Gender (female vs. male)** | 38.3 | **p=0.02\*** | 0.13 | p=0.39 | 1.44 | **p=0.006\*\*** |
|  | **Mother's Education (Beyond HS vs. not beyond HS)** | -11.9 | p=0.73 | -0.03 | p=0.93 | 0.20 | p=0.86 |
|  |  | **R<sup>2</sup>=0.29** |  | **R<sup>2</sup>=0.09** |  | **R<sup>2</sup>=0.37** |  |

Significance levels: \*p≤0.05, \*\*p≤0.01, \*\*\*p≤0.001
TRAP= traffic-related air pollution
ECAT= elemental carbon attributable to traffic

#### Beta diversity

The Bray-Curtis dissimilarity measure figures did not show a distinction in bacterial microbiota in sputum between TRAP exposure group, asthma status group, or gender (Fig. 5). However, the Adonis test indicated that 6% of the variance in the distance matrix between the sputum of asthmatics and non-asthmatics could be attributed to asthma status (p=0.04). The Adonis test also indicated that 6% of the variance in the distance matrix between the sputum of male and female participants could be attributed to gender (p=0.04). The homogeneity of dispersion test showed no significant difference in the variances of the sputum samples from each asthma status group (p=0.88) or from each gender (p=0.46) indicating no confounding effect from a difference in variances. The Adonis test did not indicate that the variance in sputum between TRAP exposure groups could be attributed to TRAP exposure (p=0.9).

**Fig. 5-.**
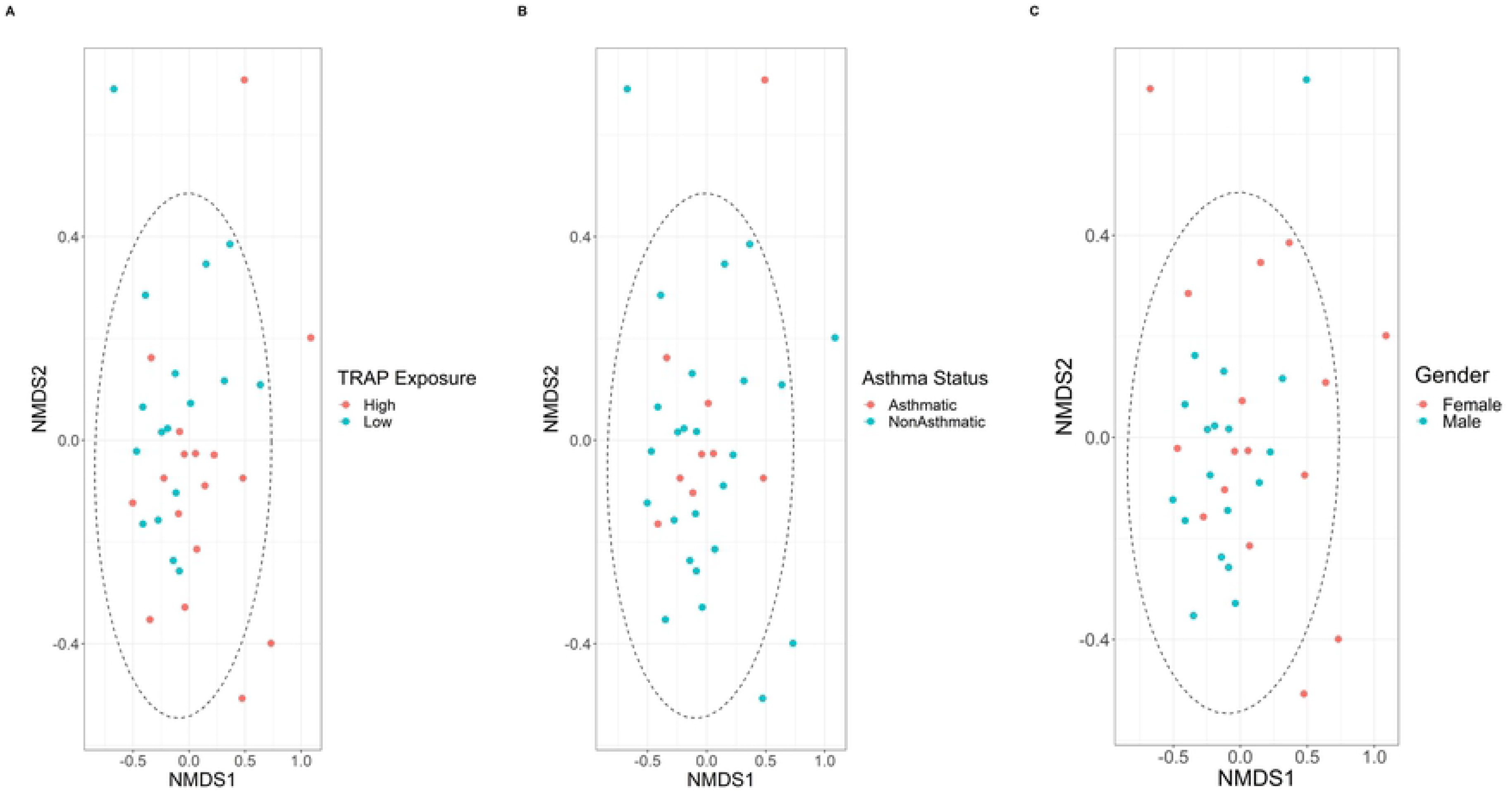
Non-metric multidimensional scaling (NMDS) of Bray-Curtis distances of bacteria in sputum microbiota from each TRAP exposure group, asthma status group, and gender.

#### Relative Abundance

According to the xdc.sevsample test, the distribution of major phyla did not differ in sputum between each TRAP exposure group nor each gender (p=0.43 and p=1.0, respectively). However, a significant difference in major phyla was found between asthma status groups (p≤0.001). These results are in agreement with the relative abundance bar plots (Fig. 6 and S5 Fig). The relative abundance of phyla in sputum does not appear to differ between TRAP exposure group nor gender, but asthmatics appear to have more *Bacteroidetes* and fewer *Proteobacteria*.

**Fig. 6-.**
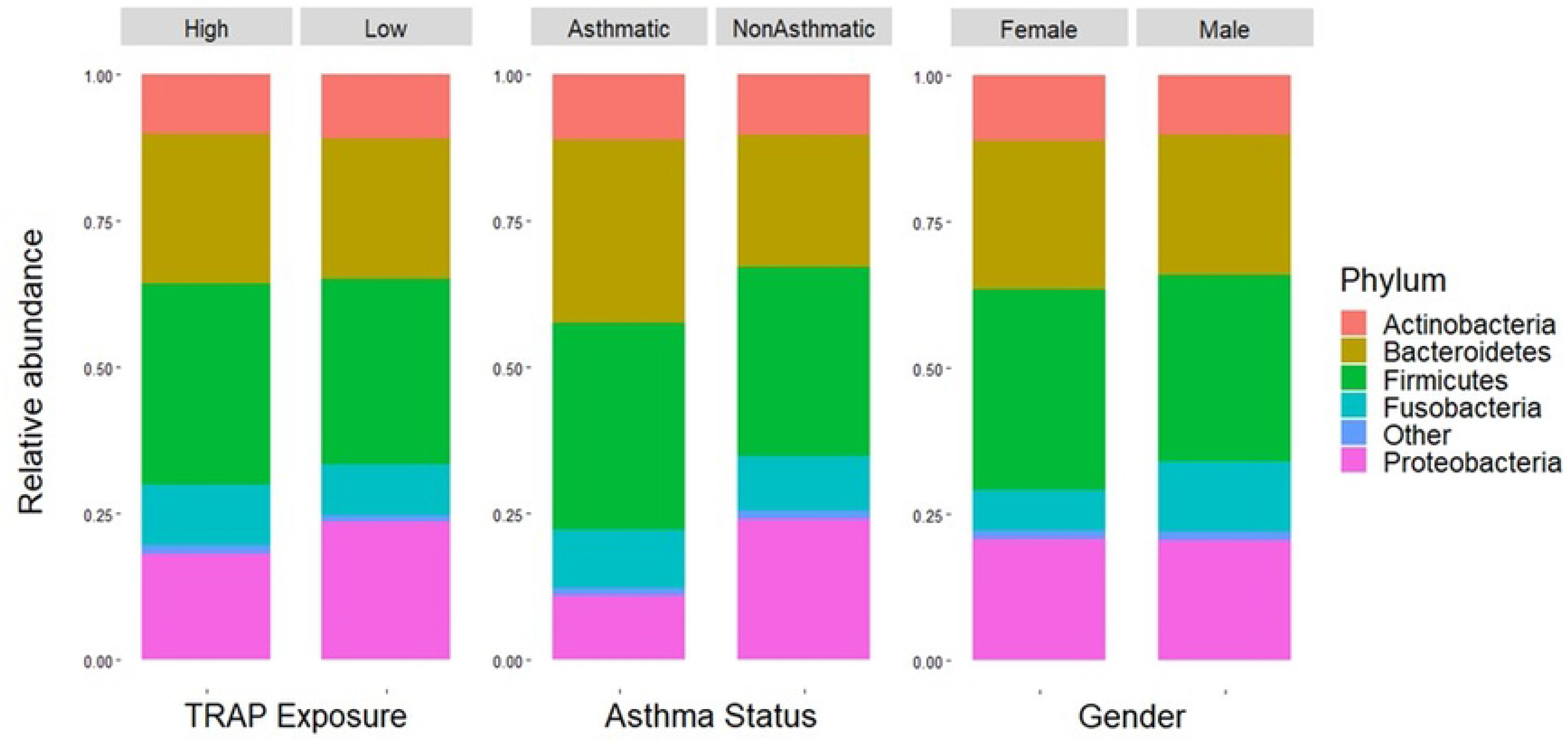
Bar plots showing the relative abundance of bacterial phyla in sputum across (A) TRAP exposure groups, (B) asthma status groups, and (C) genders.

#### Differential Abundance

While the overall microbial community composition did not appear to differ between TRAP exposure groups, negative binomial regression was able to identify several individual ASVs with a log2 fold-change greater than 2 and FDR p≤0.05 according to TRAP exposure groups (high vs. low) (Fig. 7). *Fusobacterium nucleatum* had a log2 fold change of 25 (FDR p≤0.001). Two other ASVs in the *Fusobacterium* genus had FDR p-values ≤0.001, with log2 fold-changes of 24 and 9, and one ASV in the genus *Atopobium* (family *Atopobiaceae*) had a log2 fold-change of 7 (FDR p=0.02) across the TRAP exposure groups. While the FDR p-values for the ASVs in the family *Prevotellaceae* were not <0.05, the log2 fold-changes were more consistently >2 for individual ASVs in the family.

**Fig. 7-.**
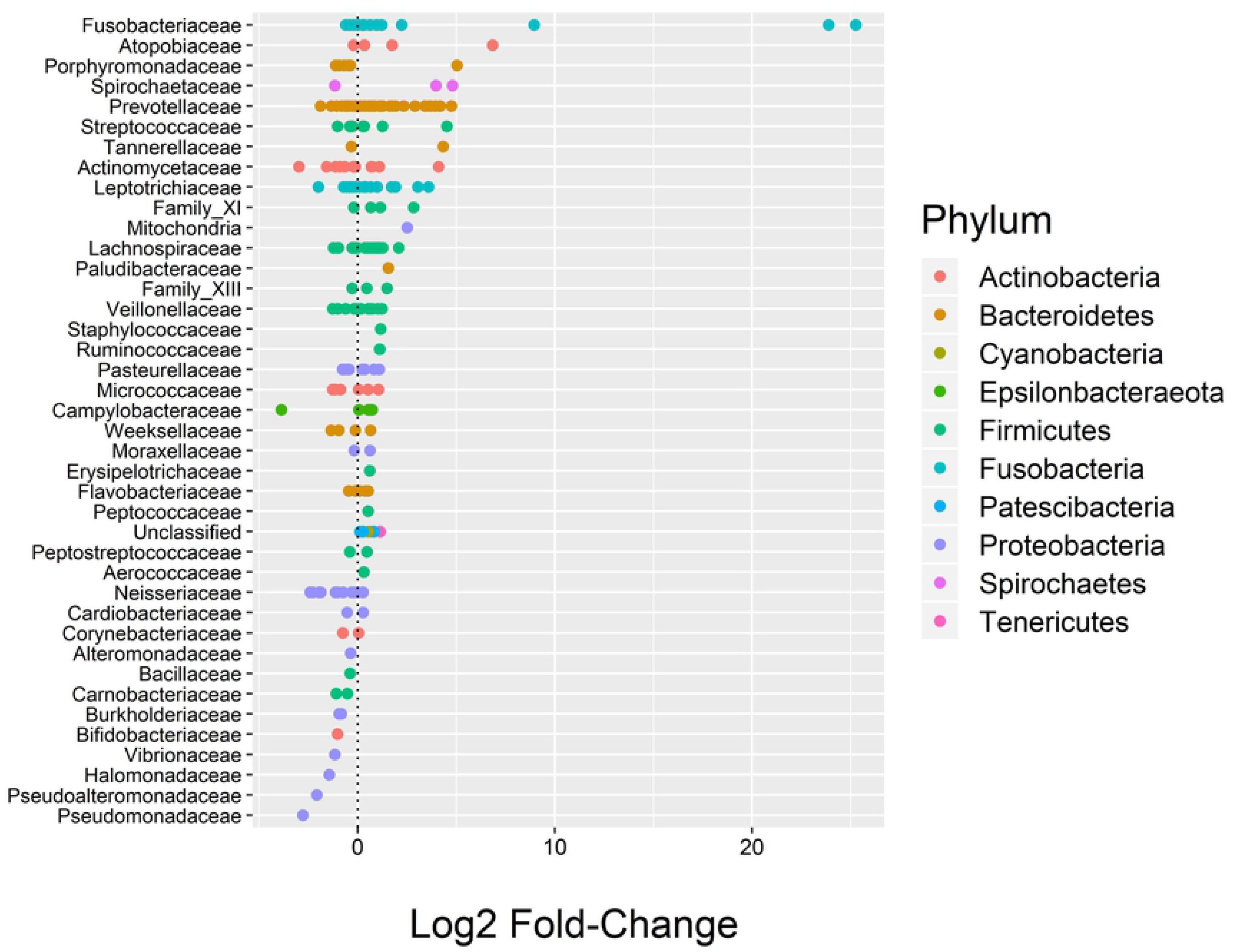
DESeq2 results showing the log2 fold-change values (x-axis) of bacteria in sputum between TRAP exposure groups. Each line on the y-axis indicates the family, each point represents an individual ASV within that family, and the color of the point indicates the phylum.

We also identified several ASVs there were differentially abundant in sputum across gender (female vs. male) and two ASVs that were differentially abundant in sputum across asthma status groups (asthmatic vs. non-asthmatic) (S6 and 7 Figs). Across gender, there were two ASVs in *Fusobacterium* with log2 fold-changes of 24 and 25 (FDR p≤0.001), one ASV in the genus *Campylobacter* (family *Campylobacteraceae*) with a log2 fold-change of −8 (FDR p=0.01), and one ASV in the genus *Prevotella* (family *Prevotellaceae*) with a log2 fold-change of −7 (FDR p=0.03). Across asthma status, there was one ASV identified as *Prevotella salivae* with a log2 fold-change of −3 (FDR p=0.008), and one ASV in the genus *Bacillus* (family *Bacillaceae*) with a log2 fold-change of 8 (FDR p=0.05).

### Overall Bacterial Load

We compared the overall bacterial load in sputum, as measured by qPCR with universal bacterial primers, between TRAP exposure groups, between asthmatics and non-asthmatics, and between genders using Wilcoxon rank sum test. No significant difference in the overall bacterial load was observed between high and low TRAP exposure groups (median high TRAP= 6.9 x10^5^ bacterial genome copies per mL of sputum, median low TRAP=3.6 x 10^5^ bacterial genome copies per mL of sputum; p=0.43) (Fig. 8). However, there was a higher bacterial load in the sputum of asthmatic participants than in non-asthmatic participants (median asthmatic=1.3 x 10^6^ bacterial genome copies per mL of sputum, median non-asthmatic=3.6 x 10^5^ bacterial genome copies per mL of sputum; p=0.07) and in the sputum of male than in female participants (median male=9.4 x 10^5^ bacterial genome copies per mL of sputum, median female=1.7 x 10^5^ bacterial genome copies per mL of sputum; p=0.006) (Fig. 8). We also found a higher bacterial load in the saliva of asthmatics (median asthmatic=4.0 x 10^6^ 10^5^ bacterial genome copies per mL of saliva, median non-asthmatic=1.3 x 10^6^ 10^5^ bacterial genome copies per mL of saliva; p=0.05). There were no differences in the bacterial load in saliva between TRAP exposure groups (p=0.26) nor between genders (p=0.65).

**Fig. 8-.**
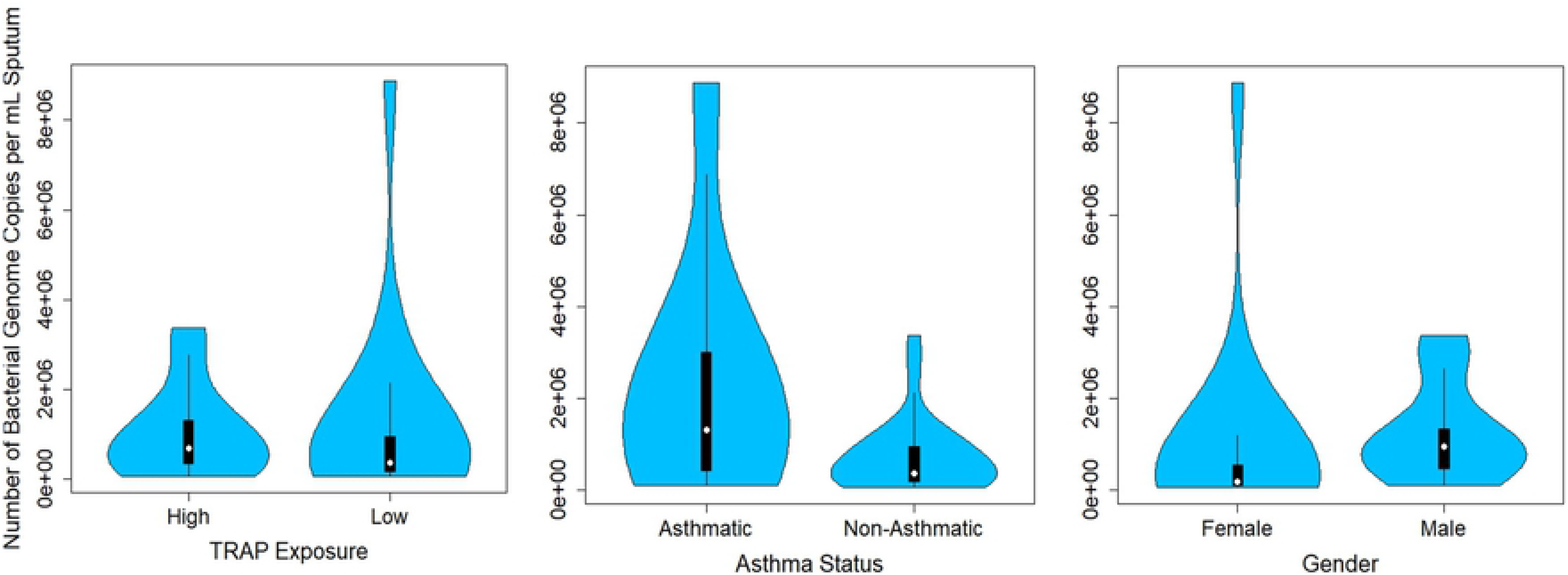
Violin plot comparing the total bacterial load, as measured by qPCR, in the sputum of each TRAP exposure group, each asthma status group, and each gender.

### Traffic Pollution and the Respiratory Fungal Microbiota (Mycobiome)

We compared the sputum mycobiome of high and low TRAP-exposed participants, as well as by gender and asthma status. The median ECAT for the high exposure group was 0.40 mg/m^3^ and the median for the low exposure group was 0.29 mg/m^3^. There was an overlap between confidence intervals for all three alpha diversity measures when comparing TRAP exposure groups, genders, and asthma status groups (S1 Table). There also were no significant differences in beta diversity (Bray-Curtis) between asthma status groups (p=0.35), between TRAP exposure groups (p=0.94), nor between genders (p=0.23), using the Adonis test. The xdc.sevsample test showed a difference in the distribution of major fungal classes between asthma status groups (p=0.009), and between gender (p<0.001), but not between TRAP exposure groups (p=1.0). It should be noted, however, that due to overall low fungal abundance in sputum, we had a very small sample size.

## Discussion

### Distinction Between Sputum and Saliva

The most abundant bacterial phyla in both sputum and saliva samples were *Actinobacteria, Firmicutes, Bacteroidetes*, and *Proteobacteria*. The major phyla identified in the sputum and saliva samples are consistent with those found in previous studies [42, 76, 77]. The Bray-Curtis dissimilarity measures also showed a distinction between the two sample types for bacteria. The saliva and sputum samples clustered closely, but there was still a noticeable distinction between the two. The Bray-Curtis and Jaccard NMDS plots showed separation between sample type, but the UniFrac NMDS plot did not show separation between sample type. This suggests that while the abundance and presence/absence measures show separation between sample type, sputum and saliva do not appear to be different when considering the phylogenetic distances. These results are consistent with the alpha diversity analyses, where there was a significant difference between sample type in Shannon diversity and number of observed ASVs, but not phylogenetic diversity. The alpha and beta diversity results make sense when considering that while the growth condition in the lungs and the oral cavity are quite different, the migration of microbes into the airways occurs primarily through microaspiration of saliva [2]. Additionally, the differences in alpha diversity and beta diversity between sample type indicate that the sputum was not entirely contaminated by the oral microbiome.

### Bacteriome Differences in TRAP Exposure Groups

As previous studies have shown that asthmatics tend to have greater alpha diversity in their respiratory tracts than non-asthmatics, and exposure to TRAP is associated with a higher incidence of asthma, asthma status was examined as a potential confounder [33]. We compared the diversity indices between asthmatics and non-asthmatics. While the mean number of observed ASVs, Shannon diversity, and phylogenetic diversity were slightly higher in asthmatics than in non-asthmatics, the confidence intervals of asthmatics and non-asthmatics overlapped in each alpha diversity measure. In contrast, the confidence intervals of the high and low TRAP-exposure groups did not overlap in the mean number of ASVs nor the mean phylogenetic diversity. These results are consistent with the Wilcoxon rank sum test p-values and the diversity measure box plots. Thus, asthma status does not appear to have a significant impact on our results of overall diversity comparisons between high and low TRAP exposure groups. However, as there were far fewer asthmatics than non-asthmatics included in the study, it is difficult to make a clear determination of differences in relative abundance between the asthma status groups. We also observed greater alpha diversity indices in the sputum of females than in males. To our knowledge, there have been no previous studies that have documented a significant difference in the bacterial diversity of the lower respiratory tract between genders [6]. However, there have been studies of the gut and skin microbiome that have documented differences in the microbial communities between genders, with higher levels of specific taxa found in the gut of males, and overall higher bacterial diversity found on the skin of males [78, 79]. In contrast to sputum, we did not observe any significant differences in the alpha diversity indices of saliva between TRAP exposure groups, gender, or asthma status groups. This further supports the notion that the bacterial microbiome in the sputum samples is distinct from that in the saliva samples.

The results of the multiple regression model were consistent with the results of the univariate analysis (Table 2). Neither asthma status nor mother’s education were significant predictors of alpha diversity indices, further supporting that asthma status did not have a significant impact on our results of the TRAP exposure group comparisons. The results of the regression model with TRAP as a categorical variable are consistent with the Wilcoxon rank sum test results comparing diversity indices between exposure groups. The results of the regression model with TRAP as a continuous variable (ECAT) further strengthen the notion that TRAP exposure increases bacterial diversity in the lower respiratory tract. The models for phylogenetic diversity had the highest R^2^ values and the models for Shannon diversity had the lowest R^2^ values.

TRAP exposure affects the body in several ways that may be impacting the bacterial diversity in the lower respiratory tract. The bacterial communities in the airways are believed to originate primarily from microaspiration of saliva and are maintained through defense mechanisms, including mucociliary clearance and immune responses [42]. There is evidence that exposure to NO_2_, a component of TRAP, impairs mucociliary activity, which would increase the number of microbes that remain in the airways [80, 81]. Additionally, the PM component of TRAP has been shown to increase adhesion of bacteria to human airway epithelial cells [9], promote an airway inflammatory response [25-28], and reduce the production of antimicrobial peptides [29]. It is unclear which of these mechanisms contributes to the increase in diversity and should be an area of focus in future research.

Lower bacterial diversity is usually associated with diseased states in parts of the body, such as the gut [82-84]. However, in the lungs, greater bacterial diversity has been associated with a diseased state [33]. It is well-understood in microbial ecology that a diverse population protects against biological invasions and is able to use resources more efficiently [85-87]. Biodiversity acts as insurance to maintain a functioning ecosystem under abiotic perturbations and anthropogenic disturbances [88]. As TRAP has been shown to increase the adhesion of bacteria to respiratory tract epithelial cells, it is possible that the alteration in growth conditions in the TRAP-exposed respiratory tract could promote the development of a more diverse bacterial community. Additionally, this change in microbial communities may elicit a local immune response, or even impact immune system development in children. Future studies should examine the relationship between bacterial diversity in the lungs and respiratory health.

There was not a significant difference in beta diversity between the TRAP exposure groups. This indicates that while alpha diversity in sputum is associated with traffic pollution exposure within the range of observed values, the microbial composition between the high and low exposure groups is not significantly different. The relative abundance of bacterial phyla in sputum also did not appear to differ between TRAP exposure group or gender, but asthmatics appeared to have more *Bacteroidetes* and fewer *Proteobacteria*. Inconsistent with our results, previous studies have documented higher levels of *Proteobacteria* in asthmatics compared to non-asthmatics [33, 89]. Additionally, the overall bacterial load was significantly greater in asthmatic than non-asthmatic participants. This result is consistent with previous research that has shown higher bacterial loads in asthmatics [34]. However, it should be noted that we had fewer non-asthmatic than asthmatic participants included in this study.

While the overall microbial community composition did not appear to differ between TRAP exposure groups, negative binomial regression was able to identify several differentially abundant individual ASVs between TRAP exposure groups, including *Fusobacterium nucleatum* and one ASV in the genus *Atopobium*. ASVs in *Prevotella* did not have a statistically significant FDR p-value, but consistently had >2 log2 fold-changes. *Fusobacterium* is associated with several human diseases, most commonly periodontal and oropharyngeal infections, produces a potent endotoxin, and is known to assist in the development of biofilms [90-92]. Biofilms are known to form in the lungs of cystic fibrosis patients, so it is possible that the biofilm property of *Fusobacterium* may be relevant to respiratory health [93]. *Atopobium* is associated with bacterial vaginosis [94]. *Prevotella* is associated with anaerobic lower respiratory tract infections [95]. While we cannot meaningfully remark on the biological significance of these findings, these may be taxa of interest in future investigations regarding TRAP exposure and the respiratory microbiome. We also identified differentially abundant ASVs across gender and asthma status. Across gender, there were two differentially abundant ASVs in *Fusobacterium*, one ASV in *Campylobacter*, and one ASV in *Prevotella*. A study on the gut microbiome also identified higher levels of *Prevotella* in males than in females [78]. Across asthma status, there was one differentially abundant ASV identified as *Prevotella salivae*, and one ASV in *Bacillus*. It has been proposed that a higher abundance of *Prevotella* in the lungs may increase airway inflammation [96].

### Traffic Pollution and the Respiratory Mycobiome

Our pilot study results did not indicate a difference between the mycobiomes of sputum in the high and low TRAP-exposed participants. This could be because we had a small sample size and the samples that did amplify had very low abundance. Additionally, fungi do not proliferate to the same extent as bacteria in the lower respiratory tract. When comparing bacteriomes, we had enough samples to conduct a linear regression to determine the association between alpha diversity and TRAP exposure, both as a categorical variable (high vs. low) and continuous variable (ECAT), while adjusting for gender, asthma status, and socioeconomic status. Unfortunately, because of the small sample size we could not do the same for the mycobiome. Other sequencing methods besides marker gene analysis, such as shotgun metagenomics, may be better choices for these types of samples and should be explored further in future studies.

### Strengths and Limitations

A major strength of this study is the well-characterized TRAP exposure history of the CCAAPS cohort. Due to this characterization, we were able to examine the effect of TRAP exposure on bacterial diversity in the lower respiratory tract as both a categorical and continuous variable. However, we were limited by the small sample size. Additionally, while the data support that asthma status did not significantly impact our bacterial results, it must be noted that we had very few asthmatics compared to non-asthmatics in this study. We also did not have the specific endotypes of the asthmatic participants. Previous studies have shown that specific asthma endotypes may impact the respiratory microbiome in different ways [42].

Another strength of this study is that we conducted both 16s and ITS metagenomic sequencing and qPCR with universal bacterial and fungal primers on all samples. Therefore, we were able to normalize our sequencing results with the qPCR data instead of relying on statistical methods, such as rarefaction, to account for sequencing depth. One limitation of the qPCR method is that for both bacteria and fungi, species contain variable numbers of the amplified genomic region [56]. Therefore, the measurement of the total number of bacterial and fungal genome copies per mL of sputum may be affected by the species present in the sample.

We selected the induced sputum method over bronchoalveolar lavage because it is less invasive. Therefore, oral contamination of the sputum samples was a major concern in this study. However, one strength of this study is that our results demonstrated that the saliva and sputum samples had distinct bacterial communities, indicating that the sputum samples were not entirely contaminated by the oral microbiome. It should also be noted that the lungs have a wide range of microgeographic conditions, with a temperature gradient from ambient air temperature to body temperature in the short distance from the point of inhalation to the alveoli, so future studies may want to focus on specific locations within the lower respiratory tract.

## Conclusions

These findings indicate that exposure to TRAP in early childhood and adolescence is associated with greater diversity in the lower respiratory tract in our sample of participants. It is still unknown whether the development of asthma changes the lower respiratory tract microbiome or if an altered microbiome mediates a change in disease status. However, these results demonstrate that there may be a TRAP-exposure-related change in the lower-respiratory microbiome that is independent of asthma status. We also identified several taxa of interest for future studies, including *Fusobacterium, Atopobium*, and *Prevotella*. A major limitation of this study was the small sample size, so a larger pool of participants is needed to confirm our findings. Additionally, a study with a larger sample size could use model-based approaches for a more robust examination of the relationship of TRAP exposure and the respiratory mycobiome. Further research characterizing the human microbiome in relation to environmental exposures can lead to important new discoveries into how the body is impacted by these exposures.

## Acknowledgments

The content is solely the responsibility of the authors and does not necessarily represent the official views of the NIH. The induced sputum samples were collected with the assistance of the Schubert Research Clinic at Cincinnati Children’s Hospital Medical Center. The authors are thankful for Reshmi Indugula, who assisted with the qPCR analyses.

## List of Abbreviations

PM: particulate matter
TRAP: traffic-related air pollution
ECAT: environmental carbon attributable to traffic
LUR: land-use regression
ASV: amplicon sequence variant

## Supporting information

**S1 Fig. Bar plots showing the relative abundance of each bacterial phyla in sputum and saliva.**

**S2 Fig. NMDS (upper graphs) and MDS (lower graphs) plots using Bray-Curtis, unweighted UniFrac, and Jaccard beta diversity measurements to compare the bacteriomes in sputum and saliva**

**S3 Fig. Box plots comparing bacterial diversity indices in sputum between (A) asthma status and (B) gender, including Shannon diversity, number of observed amplicon sequence variants (ASVs), and Faith’s phylogenetic diversity.**

**S4 Fig. Box plots comparing bacterial diversity indices in saliva between (A) TRAP exposure groups, (B) asthma status groups, and (C) gender, including Shannon diversity, number of observed amplicon sequence variants (ASVs), and Faith’s phylogenetic diversity.**

**S5 Fig. Bar plots showing the relative abundance of bacterial phyla in sputum across TRAP exposure groups, asthma status groups, and genders.**

**S6 Fig. DESeq2 results showing the log2 fold-change values (x-axis) in sputum bacteriome between genders. Each line on the y-axis indicates the family, each point represents an individual ASV within that family, and the color of the point indicates the phylum.**

**S7 Fig. DESeq2 results showing the log2 fold-change values (x-axis) in sputum bacteriome between asthma status groups. Each line on the y-axis indicates the family, each point represents an individual ASV within that family, and the color of the point indicates the phylum.**

**S1 Table, Mean (95% confidence interval) and Wilcoxon rank sum test p-value for each fungal alpha diversity measure in the sputum by TRAP exposure, asthma status, and gender.**

